# *Glial swip-10* expression controls systemic mitochondrial function, oxidative stress, and neuronal viability via copper ion homeostasis

**DOI:** 10.1101/2023.12.06.570462

**Authors:** Peter Rodriguez, Vrinda Kalia, Chelsea L. Gibson, Zayna Gichi, Andre Rajoo, Carson D. Matier, Aidan T. Pezacki, Tong Xiao, Lucia Carvelli, Christopher J. Chang, Gary W. Miller, Andy V. Khamoui, Jana Boerner, Randy D. Blakely

## Abstract

Cuprous copper (Cu(I)) is an essential cofactor for enzymes supporting many cellular functions including mitochondrial respiration and suppression of oxidative stress. Neurons are particularly dependent on these pathways, with multiple neurodegenerative diseases, including Alzheimer’s disease (AD), Parkinson’s disease, associated with their dysfunction. Key features of Cu(I) contributions to neuronal health *in vivo* remain to be defined, owing largely to the complex processes involved in Cu(I) production, intracellular transport, and systemic redistribution. Here, we provide genetic and pharmacological evidence that *swip-10* is a critical determinant of systemic Cu(I) levels in *C. elegans*, with deletion leading to systemic deficits in mitochondrial respiration, production of oxidative stress, and neurodegeneration. These phenotypes can be reproduced in wild-type worms by Cu(I)-specific chelation and offset in *swip-10* mutants by growth on the Cu(I) enhancing molecule elesclomol, as well as by glial expression of wildtype *swip-10*. *MBLAC1*, the most closely related mammalian ortholog to *swip-10*, encodes for a pre-mRNA processing enzyme for H3 histone, a protein whose actions surprisingly include an enzymatic capacity to produce Cu(I) via the reduction of Cu(II). Moreover, genome-wide association studies and post-mortem molecular studies implicate reductions of *MBLAC1* expression in risk for AD with cardiovascular disease comorbidity. Consistent with these studies, we demonstrate that the deposition of β-amyloid plaques, an AD pathological hallmark, in worms engineered to express human Aβ_1-42,_ is greatly exaggerated by mutation of *swip-10*. Together, these studies identify a novel glial-expressed, and pathway for Cu(I) production that may be targeted for the treatment of AD and other neurodegenerative diseases.

**Significance Statement:** Devastating neurodegenerative diseases such as Alzheimer’s disease, and Parkinson’s disease are associated with disruptions in copper (Cu) homeostasis. Alterations in Cu(I) give rise to increased oxidative stress burden, mitochondrial and metabolic dysfunction, and can accelerate production and/or potentiate toxicity of disease-associated protein aggregates. Here, using the model system *Caenorhabditis elegans*, we establish a role for the gene *swip-10* in systemic Cu(I) homeostasis. Perturbation of this pathway in worms recapitulates biochemical, histological, and pathological features seen in human neurodegenerative disease. We reveal that these changes can be suppressed pharmacologically and arise when *swip-10* expression is eliminated from glial cells. Our work implicates *swip-10* and orthologs as key players in Cu(I) homeostasis that may be exploitable to treat multiple neurodegenerative diseases.

## Introduction

Copper (Cu) is an essential micronutrient involved in numerous fundamental aspects of cell physiology including mitochondrial respiration, ATP production, suppression of oxidative stress, redox-dependent biosynthetic pathways, and metal ion-supported cell signaling(1-3). The ability of Cu to readily accept and donate electrons are essential to the activities of Cu-dependent proteins(4). Among the most well studied roles for Cu are the detoxification of reactive oxygen species (ROS) and support for mitochondrial electron transport chain (ETC) function via activation of superoxide dismutase (SOD) and cytochrome c oxidase (CytC), respectively(5). In turn, mitochondrial function, and ROS buffering are key determinants of neuronal health and signaling(6). Indeed, a large amount of dietary Cu ultimately is stored and used in the brain (∼3%), second only to the liver in Cu concentrations(7). Menke’s Disease and Wilson’s Disease, genetic disorders characterized by systemic diminished and excess Cu concentrations, respectively, result in neurodegeneration(8). Altered CNS Cu homeostasis has also been linked to more common neurodegenerative diseases (NDDs), including Alzheimer’s disease (AD) and Parkinson’s disease (PD)(8). Cu exists in biological systems primarily in one of two ionic forms, specifically cuprous Cu (Cu(I)) and cupric Cu (Cu(II)). Cu(I) and Cu(II) are handled by different chaperones and transporters, and the interconversion between the two differentially supports a host of enzymatic reactions. How a proper balance of these Cu forms is achieved continues to be an active area of investigation. Recent studies using the *Saccharomyces cerevisiae* model have revealed a vital role of histone H3:H4 complexes in mitochondrial function and suppression of oxidative stress. Namely, H3:H4 complexes facilitate Cu(I) production from Cu(II), an enzymatic reductase activity independent of the role of these proteins in DNA compaction and gene regulation(9), in keeping with the key role played by Cu(I) in mitochondrial electron transport by cytochrome C oxidase (CytC) and superoxide dismutase (SOD)(5). Whether this pathway plays a role in neuronal health and signaling has yet to be established, though the metabolic and growth defects of a yeast model of Friedreich’s ataxia can be rescued through manipulation of H3 Cu reductase activity(10).

Our lab has utilized the genetic model system *Caenorhabditis elegans* to elucidate novel genetic pathways that support signaling and health of multiple classes of neurons including those secreting dopamine (DA), acetylcholine (ACh), and glutamate (Glu)(11-16). For example, we demonstrated that worms labeled through DA neuron-specific green fluorescent protein (GFP) expression are damaged, like mammalian DA neurons, by brief incubations with the neurotoxin 6-OHDA that is prevented in animals lacking the deletion of the gene encoding the presynaptic DA transporter (*dat-1*), a known mediator of DA neuron-specific uptake of the toxin. Subsequently, we found that *dat-1* deletion leads to rapid paralysis when mutant animals are placed in water(17), a phenotype termed Swimming Induced Paralysis (Swip). The robust and highly reproducible nature of the Swip phenotype prompted us to adopt it for a forward genetic screen to identify molecules required to support of the function and health of DA neurons(18). Importantly, this screen identified multiple lines with mutations in *dat-1*, whose Swip phenotype could be reversed through pharmacological or genetic elimination of presynaptic DA stores and postsynaptic DA receptors(12, 18, 19).

Among the genes identified in our Swip screen, swip-*10*, a previously unstudied gene in *C. elegans*, was found to act cell non-autonomously via its expression in glial cells to limit DA signaling(18). Our functional studies of *swip-10* mutants revealed Glu-dependent hyperexcitability of DA neurons that triggers excess DA release and suppression of movement by the DA receptor DOP-3(18, 19). Excess Glu signaling onto mammalian DA neurons results in DA neuron degeneration and has been proposed to be a contributor to PD(20, 21). In this regard, our subsequent analysis revealed a premature death of DA neurons in *swip-10* mutants that could be attenuated by genetic elimination of Glu receptors(15). Interestingly, we also observed that *swip-10* effects extended beyond DA neurons, as degeneration could be observed in other neurons, specifically ones ensheathed by glia whereas those lacking ensheathment appear unaffected. These findings reinforce a critical role of glia in supporting neuronal health and signaling(15), and Glu-dependent neurodegeneration that can arise from disrupted glial-neuron interactions(22). Lastly, we found elevations of whole body oxidative stress as reported by the reporter *gst-4*:GFP, suggesting that, as seen with the systemic control of proteostasis(23, 24), glial cells play an important role in the worm in limiting body-wide ROS production.

Sequence alignment of *swip-10* to vertebrate genomes revealed strongest homology in humans to the human gene *MBLAC1*, driven by their shared Metallo β-lactamase Domain (MBD)(14). Conspicuously, both mutations recovered in our screen reside in the *swip-10* MBD, suggesting that a causal role for disruptions in the enzymatic function of this domain in glial support of neural activity and health in worms, and possibly in humans with diminished expression of *Mblac1*. Subsequently, Pettinati and colleagues demonstrated that the MBD of *MBLAC1* encodes a 3’ endonuclease activity that processes pre-mRNA of replication dependent (RD), H3 histone proteins(25). Although these authors clearly demonstrated a role for MBLAC1 in cell cycle progression in adenovirus transformed kidney cells *in vitro*, an *in vivo* role for the protein in non-transformed cells was not established. Indeed, both *Mblac1*^-/-^ mice(26) as well as *swip-10* mutant worms show no gross physical abnormalities, developmental defects, or lifespan alterations that would suggest a key, non-redundant contribution of *swip-10* to cell cycle regulation. Importantly, RD histones are actively transcribed in post-mitotic cells(27), including neurons, where they act to regulate gene expression(27, 28). Following the recent findings in yeast that H3:H4 histone complexes possess the capacity to reduce Cu(II) to Cu(I), we pursued the ability of *swip-10* to regulate Cu ion homeostasis and thereby promote neuronal viability and more broadly, systemic mitochondrial function and suppression of oxidative stress. Moreover, due to a genome wide association finding of *MBLAC1* as a risk factor for AD with cardiovascular comorbidity (AD-CVD)(29), accompanied by findings of reduced cortical expression of *MBLAC1*, we asked whether *swip-10* impacts pathological features of AD when modeled in the worm.

## Results

### Systemic Cu(I) homeostasis is supported by expression of *swip-10*

To determine whether loss of *swip-10* expression perturbs Cu(I) homeostasis *in vivo*, we took advantage of the recently developed, Cu(I)-specific fluorescent probe, Copper Fluor-4 (CF4)(30). As previously shown for CF4 staining in WT (N2) nematodes(31), Cu(I) accumulates systemically in lysosome-like intestinal granules. Staining of *swip-10* mutants revealed significantly fewer and noticeably larger CF4 labeled granules vs WT animals (**Fig. 1A, B**) whereas a CF-4 analog lacking Cu(I) selectivity fails to distinguish between genotypes (CF4-Control, **Fig. 1C,D**). As a further test of the *in vivo* Cu(I) specificity of CF4, we grew worms on plates containing the Cu(I) chelator bathocuproinedisulfonic acid (BCS), finding CF4 signal to be reduced to levels detected with CF4-Control (**Supp. Fig. 1**). Consistent with these findings, prior work has shown that granular staining is also depleted by loss of CUA-1, the *C. elegans* ortholog of the mammalian subcellular Cu(I) transporter ATP7A/B(31).

**Figure 1.**
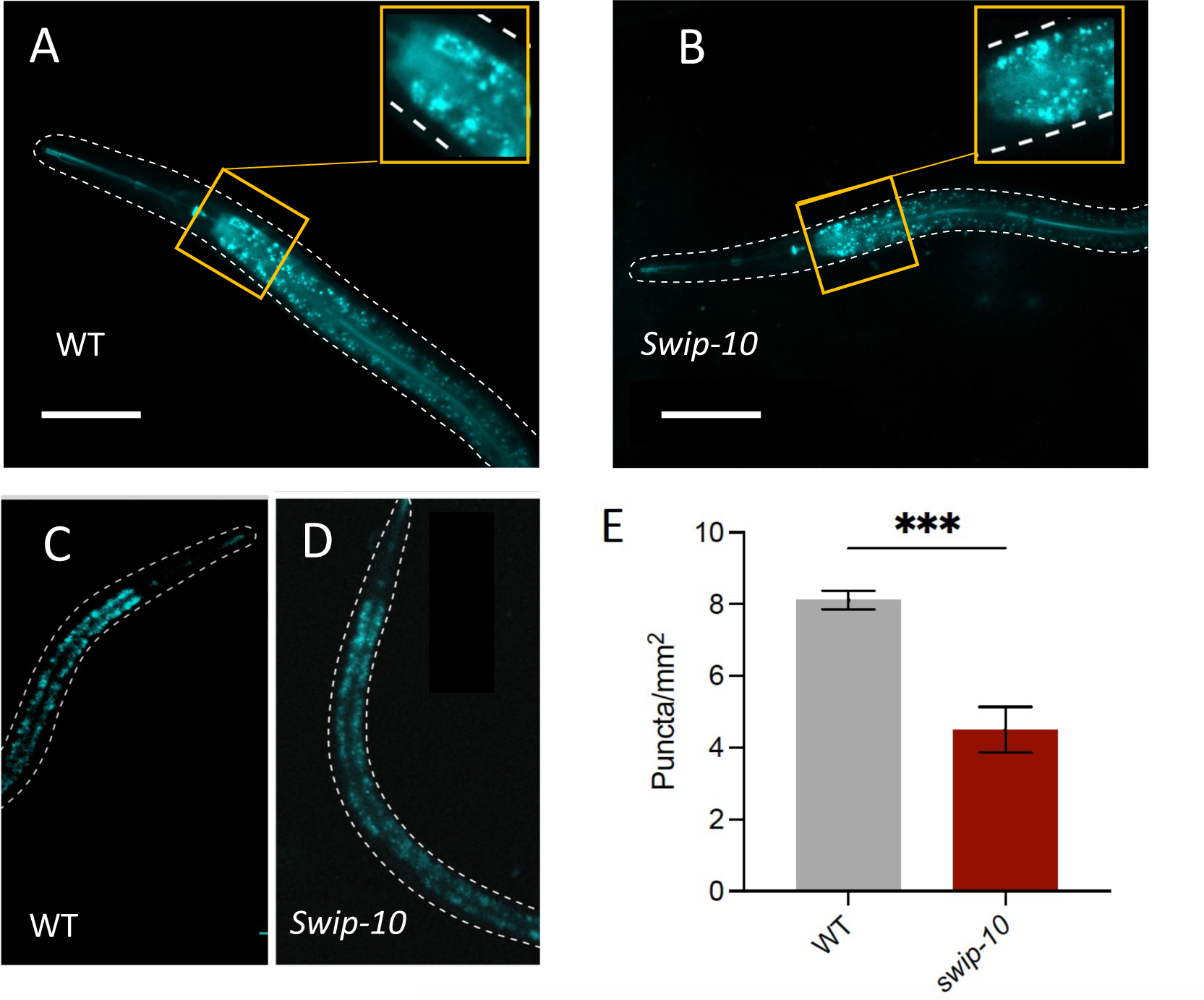
Global Cu(I) levels are sustained by expression of *swip-10*. A,B) Representative images of CF4 treated worms obtained via 20x confocal microscopy, using excitation laser at 540nm. C,D) Representative images of control CF4 treated worms obtained via 20x confocal microscopy, using excitation laser at 540nm. C) WT animals, D) *swip-10* animals. E) Quantitation of puncta observed in CF4 treated animals. Statistical analyses performed using Student’s t-test. \*\*\**p* ≤0.001. Scale bars equal to 50 microns.

### Loss of *Swip-10* impairs systemic mitochondrial respiration, ATP production and leads to oxidative stress *in vivo*

The vital role played by Cu(I) in sustaining mitochondrial function and suppressing oxidative stress led us to explore whether these capacities are impacted in *swip-10* mutants. To this end, we first measured basal O_2_ consumption in living worms using an Oroboros Oxygraph 2k respirometer. These studies revealed a nearly 50% reduction in O_2_ consumption rate (OCR) in *swip-10* mutants compared to WT animals, consistent with a significant perturbation of mitochondrial energy production (**Fig. 2A**). Consistent with these results, follow-up studies revealed significantly reduced ATP levels in *swip-10* mutants (**Fig. 2B**). Although a sensitive platform, the Oroboros system limited our ability to detect changes in OCR by pathways independent of mitochondrial oxidative phosphorylation (OXPHOS) and to assess maximal capacity for mitochondrial respiration. We therefore assessed OCR using a Seahorse XF96 respirometer which provides a capacity to add drugs sequentially to the same sample (**Fig. 2C**), such as mitochondrial uncouplers and electron transport chain inhibitors(32). Here, we observed a comparable and significant reduction in basal OCR in *swip-10* mutants as observed with the Oroboros platform (**Fig. 2D**). When next we added the drug carbonyl cyanide-4 (trifluoromethoxy) phenylhydrazone (FCCP) to uncouple electron transport from ATP production, and thereby drive maximal OCR, we observed that *swip-10* mutants reached a similar OCR as WT animals (**Fig. 2E**), indicative of a perturbation of mitochondrial function in *swip-10* mutants at or before complex IV, versus a change in mitochondrial number or capacity. Upon addition of sodium azide to inhibit complex IV, which stops all mitochondrial oxygen consumption, WT and *swip-10* OCR drops significantly to reveal the level of non-mitochondrial OCR capacity, which displayed no difference between WT and *swip-10* (**Fig. 2F**). Overall, these findings are consistent with reduced basal OCR in *swip-10* mutants as arising from diminished coupling of electron transport to ATP production.

**Figure 2.**
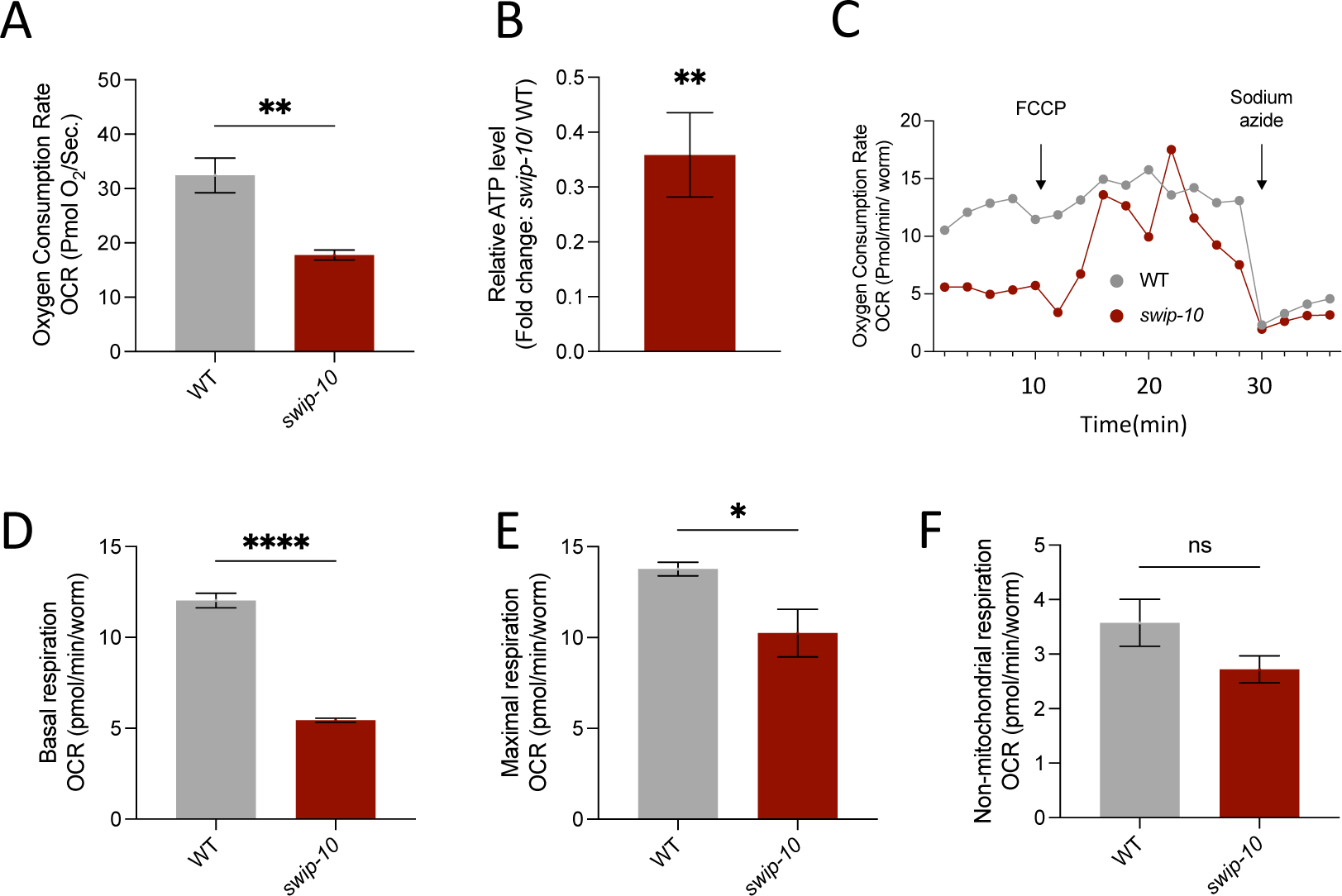
*Swip-10* mutation impairs mitochondrial activity. A) Basal oxygen consumption rate (OCR) from high-resolution respirometry with the Oroboros Oxygraph 2K. B) Quantification of ATP levels by luminometry. Data shown as average fold change of *swip-10* levels relative to N2. A Student’s t-test was used to determine significance comparing raw values (*swip-10*/N2). \**p≤* 0.05. C-F) OCR measured by Seahorse Extracellular Flux analysis. C) Representative trace recording from Seahorse experiments. X-axis displays each measurement period where probe is inserted into well and when the different drug compounds are added by the instrument. Respirometer. D) Basal recordings from animals before the addition of any drug. E) Maximal respiration induced by addition of FCCP (10μM). F) Non-mitochondrial respiration measured after the addition of the mitochondrial inhibitor sodium azide.D-F were analyzed using Welch’s t-test. \**p ≤* 0.05.

Due to the use of Cu(I) by multiple enzymes involved in ROS elimination, including mitochondrial SODs(33-35), the pronounced loss of mitochondrial function in *swip-10* mutants reported above, and our prior findings demonstrating elevated expression of an oxidative stress-sensitive reporter(15), we sought direct evidence for systemic oxidative stress in these animals. To accomplish this objective, we stained worms with the ROS-sensitive fluorophore 2’,7’-dichlorofluorescin diacetate (DCFDA) (representative images in **Fig 3A, 3B**). As quantified in **Fig. 3C**, we detected significantly elevated DCFDA fluorescence in *swip-10* mutants as compared to WT animals. We next used HPLC of worm extracts to evaluate whether ROS elevations are accompanied by alterations in reduced (GSH) and/or oxidized (GSSH) forms of glutathione, as well as how these changes impact whole worm redox potential (GSSG/GSH). We found that *swip-10* mutants exhibit a significantly decreased level of GSH (**Fig. 3D**), a non-significant elevation in GSSG (**Fig. 3E**), and a significantly decreased redox potential (**Fig 3F**) as compared to WT animals.

**Figure 3.**
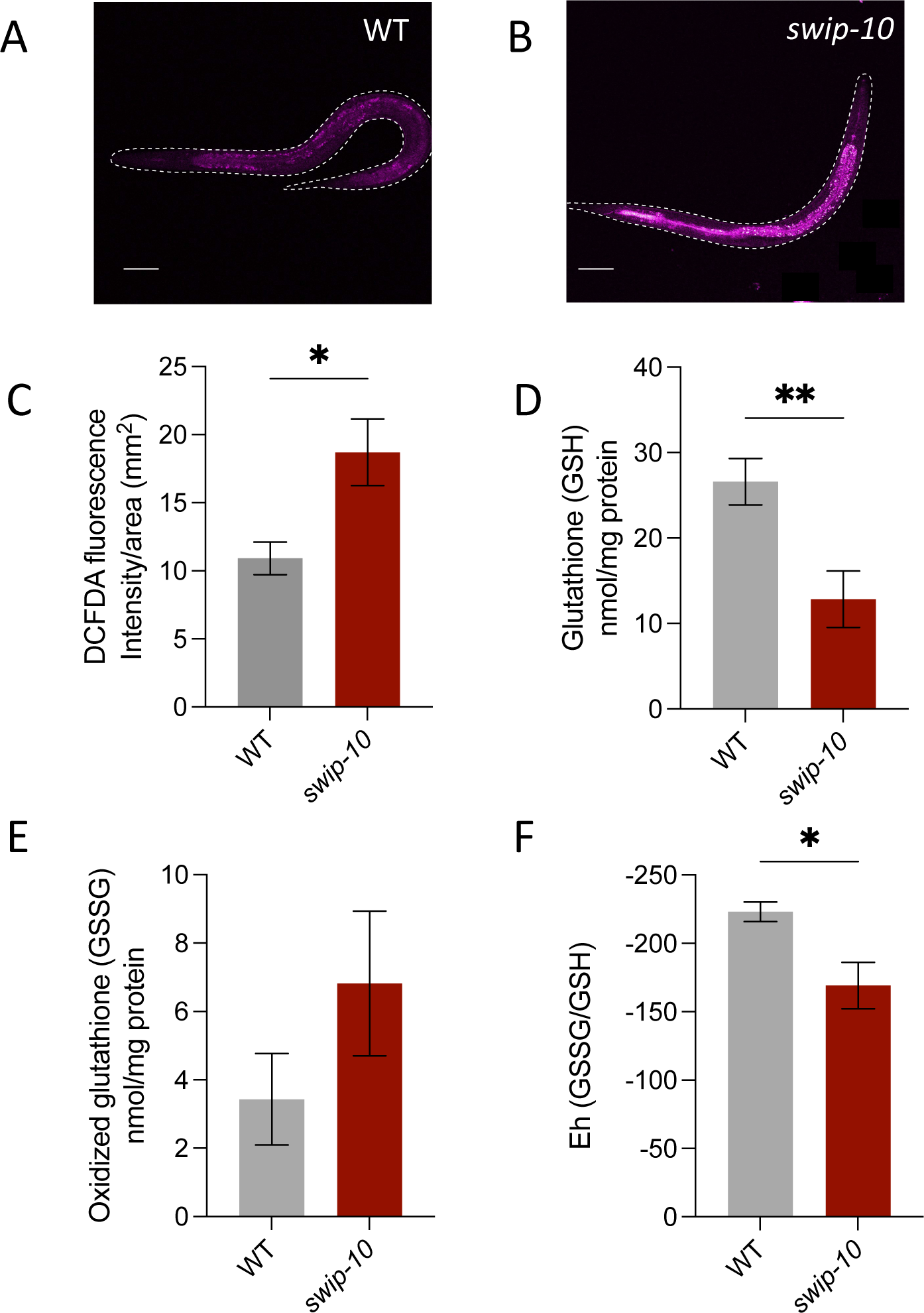
Elevated oxidative stress in *swip-10* mutants. A-B) Representative images for DCFDA obtained by confocal microscopy. 20x lens using an excitation laser of 488nm C) Quantification of fluorescent intensity of animals treated with 50μM DCFDA. Intensity average normalized to total size of animal (mm^2^). Normalized intensity/area measurements were taken from 10 animals and averaged. Analysis was performed from the average of 5 independent measurements. Statistical analysis performed using Welch’s t-test. \**p*≤0.05. 488nm. D-F) Targeted glutathione measurements using HPLC. D) Total levels of reduced glutathione (GSH). E) Total levels of oxidized glutathione (GSSG). D-E) Statistical analysis performed using Welch’s t-test. \**p*≤0.05 F) Redox potential (Eh) measured as the ratio of GSSG/GSH. Wilcoxon rank sum test performed for analysis of Eh ratio. W= 20. \**p*≤0.01. D-F) Each independent measurement was performed using approximately 500 animals. Scale bars are equal to 50μm.

### Cu(I) is both necessary and sufficient for the generation of changes in OCR, ROS, and gene expression sensitive to these alterations

If the metabolic and neurodegenerative changes observed with *swip-10* mutants derive from altered Cu(I) homeostasis, then pharmacological manipulations that restore Cu(I) levels should rescue *swip-10* phenotypes, whereas manipulations that diminish Cu(I) in WT animals should phenocopy *swip-10* phenotypes in WT animals. Indeed, both elesclomol (ES, 5µM), an organic Cu(I) chaperone(36), and CuCl_2_ (10 µM), which elevates Cu(I) in the reducing environment of the cell cytoplasm(31), fully rescued the OCR and ROS phenotypes of *swip-10* mutants (**Fig. 4A, B**). Rescue of OCR and ROS perturbations in *swip-10* animals was found to be paralleled by a normalization of expression of genes linked to Cu(I) availability, including the Cu(I) specific transporter, *chca-1* (*CTR1* ortholog), and the cytoplasmic Cu(I) chaperone *cuc-1* (*ATOX1* ortholog), which were both found to be elevated in *swip-10* mutants (**Fig 4C**). Similarly, expression of *skn-1* (ortholog of *NRF2*), *gst-4* (ortholog of *GST*), and *sod-2* (ortholog of mitochondrial *SOD2*), genes whose expression is associated with mitochondrial dysfunction and oxidative stress, were increased in *swip-10* animals, but restored to WT levels by either ES or CuCl_2_. Conversely, treatment of WT worms with the Cu(I) chelator BCS (10µM) reproduced the diminished OCR and the elevated ROS of *swip-10* mutants (**Fig. 4D and 4E**). Importantly, BCS treatment of WT animals phenocopied the DA neuron degeneration observed in *swip-10* mutants, whether assessed in aggregate (**Fig. 4F**) or broken out into individual components (**Supp. Fig. 2A-C**), whereas ES and CuCl_2_ rescued *swip-10* DA neuron degeneration (**Fig 4F,G**).

**Figure 4.**
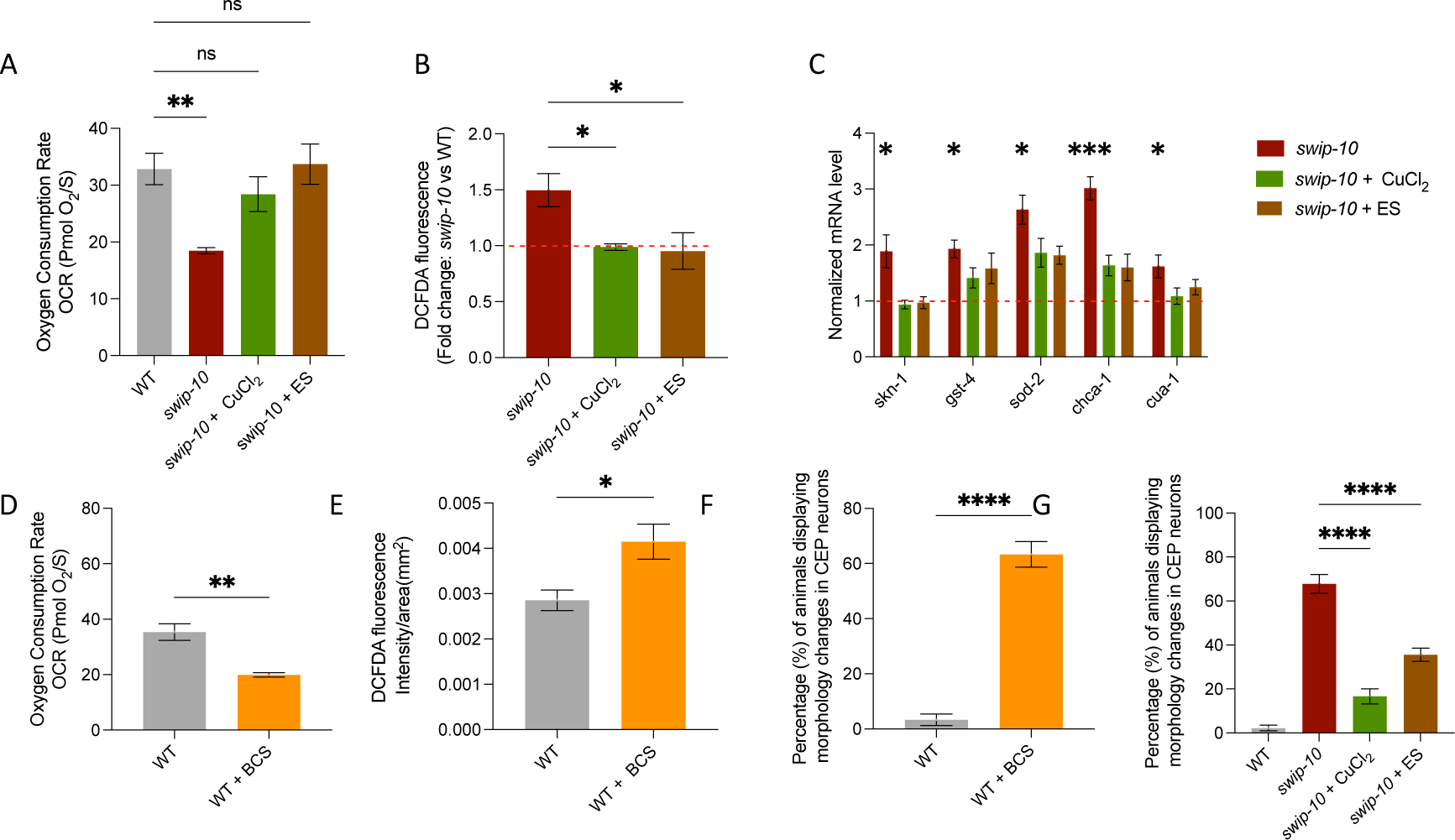
Role for Cu(I) in supporting DA neurons via homeostasis of oxidative stress and mitochondrial function. A) Basal OCR determined by Oroboros Oxygraph respirometer as an average of steady state recordings over 10 minutes. 400 animals used for recording. A,B,F) N2 animals grown on agar plates containing 10µM BCS until ready for experimentation (L4 stage). Ordinary one-way ANOVA used for analysis. \**p≤0.05.* B)Measurements of ROS levels of animals treated with DCFDA. Confocal microscopy images obtained were used to quantify relative fluorescence intensity normalized to size of each animal. Welch’s t-test used for statistical analysis. \**p* ≤ 0.05. E) Relative gene expression levels quantified via quantitative PCR reaction. All genes normalized to the housekeeping gene actin. Two technical replicates used for each measurement. N=7-8 for all experiments. Data shown as 95% confidence intervals of the mean. Dashed red line indicates the normalized value of 1 for fold change relative to wildtype (N2) animals. C,D,E)Animals either untreated or grown on agar plates containing 10µM CuCl_2_ or 5µM elesclomol. E) Basal OCR recordings (Oroboros). Ordinary one-way AVOVA used for statistical analysis. \*\**p* ≤ 0.01. B,D) Measurements of ROS levels of animals treated with DCFDA. Confocal microscopy images obtained were used to quantify relative fluorescence intensity normalized to size of each animal. Welch’s t-test used for statistical analysis. \**p* ≤ 0.05. F,G) Quantification of morphology characteristics of DA neurons. Total population percentage of animals displaying one or more characteristic of degeneration. F) Statistical analysis performed by Student’s t-test. \*\*\*\**p ≤* 0.0001. G) Statistical analysis performed by ordinary one-way ANOVA. \*\*\*\**p ≤* 0.0001.

### Glial expression of *swip-10* governs systemic Cu(I) production and changes in OCR and ROS and gene expression

Early in development, *swip-10* is expressed broadly throughout the animal, only becoming restricted to glial cells in later larval stages. Therefore, it is possible that the reductions found in Cu(I) storage granules and the alterations seen in mitochondrial function and oxidative stress in *swip-10* mutants arise from glial-independent mechanisms, despite glial requirements for *swip-10* in sustaining swimming behavior and suppressing DA neuron degeneration(14, 15). We therefore determined whether glial re-expression of WT *swip-10* selectively in glial cells supports the global phenotypes noted previously. As observed for Swip and DA neuron degeneration, driving expression of *swip-10* using pan-glial *ptr-10* promoter revealed comparable numbers of Cu(I) containing granules as detected in WT animals (**Fig. 5A**). Similarly, OCR, elevated ROS, and Cu(I) associated gene expression were also normalized **Fig. 5B-D**.

**Figure 5.**
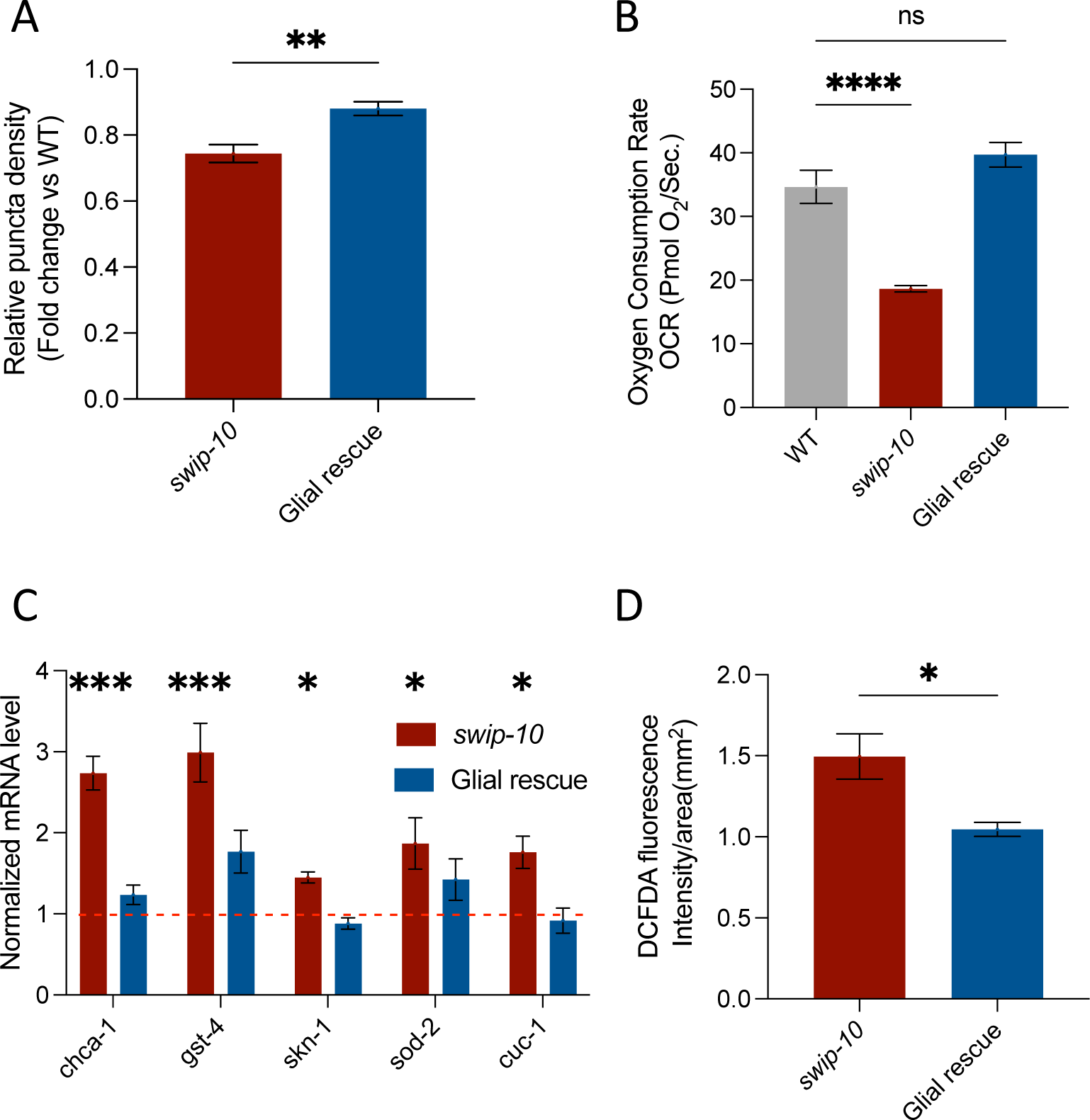
Glial expression of *swip-10* dictates global Cu(I) homeostasis, oxidative stress levels and mitochondrial function. A) Glial rescue of attenuated Cu(I) levels in *swip-10* mutants. Statistical analyses performed using Student’s T-test. \*\*\**p* ≤0.001. B) Basal OCR determined by Oroboros Oxygraph respirometer as an average of steady state recordings over 10 minutes. 400 animals used for recording. Ordinary one-way ANOVA used for analysis. \*\**p* ≤ 0.01. C) Relative gene expression levels quantified via quantitative PCR reaction. Red bars indicate changes in *swip-10* mutants, blue bars are the “glial rescue” strain. Two technical replicates used for each measurement. A minimum of 6 biological replicates were tested for all experiments. Data shown as 95% confidence intervals of the mean. Dashed red line indicates the normalized value of 1 for fold change relative to wildtype (N2) animals. D) Measurements of ROS levels of animals treated with DCFDA. Confocal microscopy images obtained were used to quantify relative fluorescence intensity normalized to size of each animal. Welch’s t-test used for statistical analysis. \**p* ≤ 0.05. Glial rescue strain contains approximately 80-90% transgenic animals expressing *swip-10* only under the pan-glial promoter (*ptr-10*).

### Metabolome perturbations of *swip-10* mutants indicate deficits in mitochondrial function and energy homeostasis

Our ability to detect changes in OCR and measures of oxidative (and ER(15)) stress in whole worms suggests that *swip-10* governs the status of multiple metabolic pathways beyond those of DA neurons To pursue this idea, we employed a global, untargeted liquid chromatography coupled high-resolution mass spectrometry (LC-HRMS)-based approach to assess metabolite alterations in *swip-10* mutants. We maximized analyte coverage of polar and non-polar metabolites by using both hydrophilic interaction LC (HILIC, under positive ionization) and C18 (under negative ionization) columns, respectively. We then employed a metabolome-wide analysis (MWA) approach, using multiple Welch’s t-tests, and a supervised dimensionality reduction approach, using partial least-squared discriminant analysis (PLS-DA), to assess the metabolic features driving the variation between N2 and *swip-10* mutant samples. These analyses revealed non-overlapping MS features for N2 and *swip-10* mutants that can be readily resolved by PLS-DA (**Fig. 6A,C**). Altogether, we found 397 features from eluates our HILIC positive column (**Fig. 6B**) and 582 features from the C18 negative column to be significantly altered comparing WT and *swip-10* samples using the MWA approach (**Fig. 6D**).

**Figure 6.**
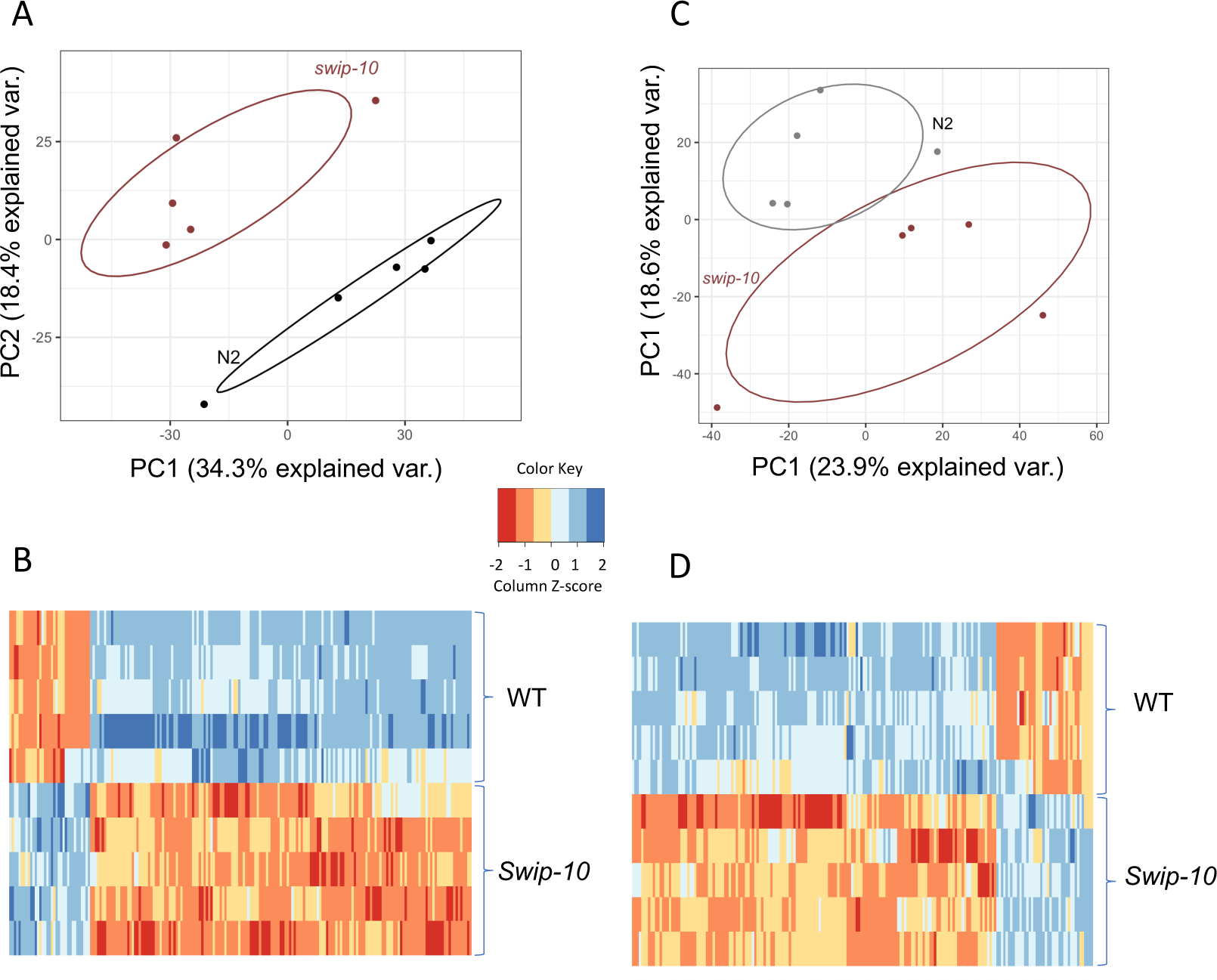
Metabolomic assessment of *swip-10* mutants. Synchronized animals were grown up to the L4 stage prior to metabolite extraction and subsequent chromatography. Raw m/z values were used for bioinformatic analyses to assess pathways altered between N2 and *swip-10* animals. A,B) PLS-DA and heatmap from HILIC positive column. C,D) PLS-DA and heatmap from C18-negative column. A,C) Statistical analyses revealed non-overlapping metabolomic signatures between genotypes. B,D) Rows represent individual biological replicate samples for each genotype. Columns represent altered metabolite changes.

To uncover metabolic pathways affected by the altered metabolic features found in *swip-10* mutants, we utilized the program mummichog(37) to assign putative identity to molecules based on the statistical likelihood of shared participation in biochemical networks. Virtually all of the pathways nominated can be directly linked to metabolic responses arising from, or contributing to, mitochondrial function and energy homeostasis including amino acid, fatty acid, sugar, steroid-hormone, and pyruvate linked metabolic pathways(38) (**Fig. 7A,B**). Conspicuously, the strongest pathway finding involves steroid biosynthesis and metabolism, processes previously found to be significantly dependent on both mitochondrial function(39) and Cu(I) homeostasis(40-42).

**Figure 7.**
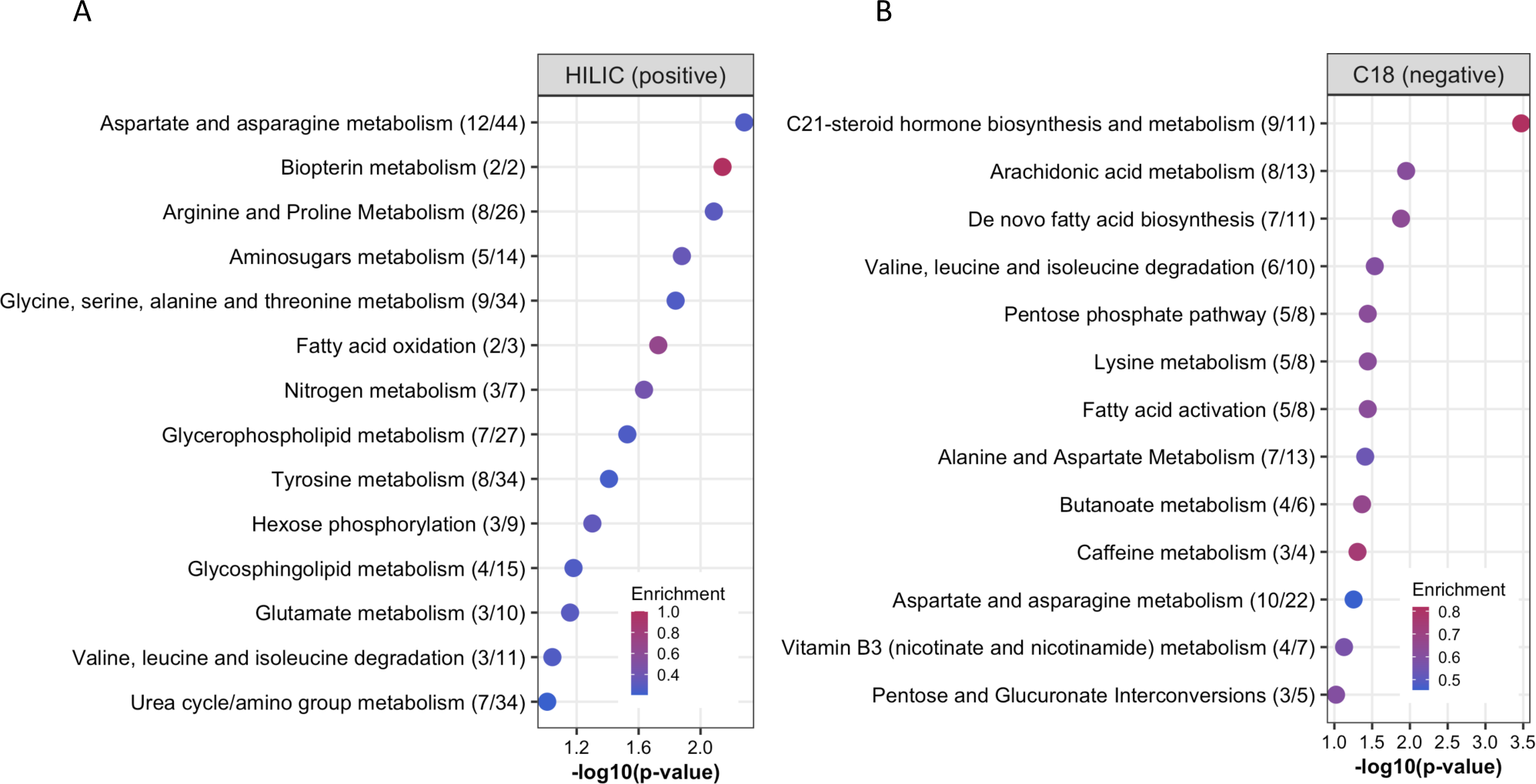
Metabolic and mitochondrial networks altered by loss of *swip-10.* Mummichog 2.0 was used for pathway assessment of altered metabolites. Enrichment scores indicates the ratio between number of metabolites altered and total pathway size. A) Metabolites measured Hilic positive column. B) Metabolites measured from C18 negative column.

Due to the high homology exhibited by the MBDs of *swip-10* and *MBLAC1*, and the report that *MBLAC1* is a risk gene for AD-CVD, we sought to determine whether a prominent pathological feature of AD, the deposition of β-amyloid plaques, is impacted by *swip-10* mutation. The GMC101 strain(43) transgenically expresses human Aβ_1-42_, the toxic, plaque forming peptide that accumulates in AD(44). We quantified plaques in the head region of GMC101 animals to avoid more posterior, intrinsic fluorescence present in non-transgenic worms, comparing levels with that found in GMC101 animals crossed onto a *swip-10* background. As shown visibly in **Fig. 8A,B** and quantified across multiple days of culture in **Fig. 8C**, we observed a striking elevation in plaque density in the context of mutant *swip-10*.

**Figure 8.**
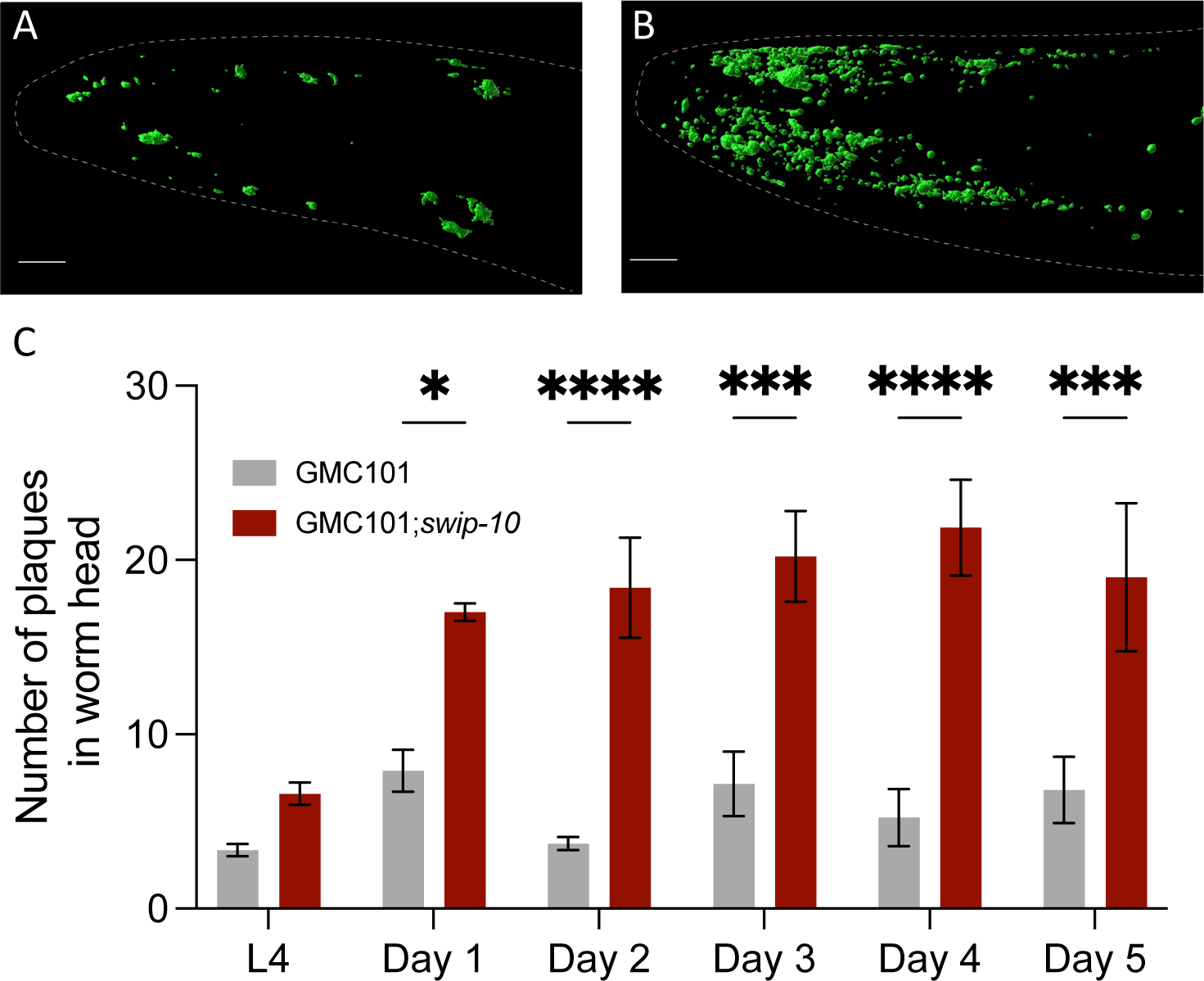
*swip-10* mutation exacerbates Aβ pathology. A,B) Representative confocal images of Aβ(1-42) expressing transgenic animals. A) GMC101 B) GMC101;*swip-10*. Number of plaques in the head of animals were manually counted from maximal projection images from the L4 stage until 4 days into adulthood. Data were analyzed using a Two-way ANOVA. Sidak’s multiple comparisons test were used to compare GMC101;*swip-10* vs. GMC101 on respective days. \*\**p* ≤ 0.01, *** p≤ 0.001. Scale bars equal to 10 µM.

## Discussion

Although Cu is critically important for metabolic functions of virtually every cell type, the energetically demanding cells of the brain are among the most sensitive to Cu dyshomeostasis(45). Not surprisingly, diseases that feature a disruption in Cu homeostasis, including Wilson’s and Menke’s disease, feature brain alterations, including AD and PD (8). Despite the critical role played by Cu ion homeostasis, the mechanisms that dictate the balance between Cu(I) and Cu(II) levels, particularly in the brain, remain ill-defined. As described above, our efforts to understand how glial *swip-10* expression non-cell autonomously regulates neuronal signaling and viability has led us to findings that the gene plays a prominent role in Cu(I) homeostasis and its role, broadly, in mitochondrial function and suppression of oxidative stress. Moreover, we show that glial cells, making up approximately 5% of total cells in the worm, support this activity and when defective can lead to neurodegeneration.

The single recognizable structural domain of *swip-10*, bearing the mutations found in our genetic screen, (18), offered us little insight as to the molecule’s function given the variety of molecules hydrolyzed by MBDs in eukaryotic proteins(46, 47). Thus, we were excited by work de-orphanizing MBLAC1, the closest mammalian SWIP-10 ortholog. This work revealed MBLAC1 to be a replication-dependent, histone pre-mRNA endonuclease. Functional studies with HEK-293 cells with diminished or absent MBLAC1 expression demonstrated alterations in cell cycle progression(25), in keeping with the well-known role played by histones in chromatin formation following DNA replication. We were puzzled, however, as to the relevance of this mechanism given the functional and neurodegenerative changes of post-mitotic DA neurons observed in *swip-10* mutants. The remarkable discovery that H3:H4 histone complexes possess an essential Cu(II) reductase activity in yeast(9) suggested an alternative hypothesis, that *swip-10* mutants may fail to support production of Cu(I), leading to deficits in mitochondrial function, oxidative stress and other vital processes that could explain both Swip and neurodegeneration. Our current study provides strong support for such a mechanism.

Initially, we compared total Cu levels in *swip-10* mutants at WT animals using inductively coupled plasma MS (ICP-MS), finding no significant changes (**Suppl. Fig. 3**). However, this approach does not distinguish Cu(I) from Cu(II) and would likely fail to detect changes in the relative ratio of these molecules since a loss of may elevate the other. We thus turned to the use of the Cu(I) specific probe, CF4, which had already been validated in worms for its ability to detect diminished Cu(I) levels arising from treatment of animals with the Cu(I) chelator BCS, also validated in our studies, as well as the loss of intestinal Cu stores due to loss of the Cu(I) transporter (CUA-1)(31). Indeed, *swip-10* mutants demonstrate a loss of Cu(I) signal, most readily quantified in counting the number of intestinal storage granules. We have not pursued the nature of these granules here, but they resemble the lysosomal-type gut granules that serve as the major storage site in worm for Zn(48). Further studies are needed to determine whether distinct granules store Zn, Cu, and other metals, whether they may discriminate Cu(I) from Cu(II), and whether their role is to eliminate high-toxic levels of Cu or can be mobilized for systemic secretion during periods of Cu insufficiency. Importantly, these granules are far from the glial cells that our rescue studies demonstrate to support Cu(I) loading, indicating that *swip-10* is important for systemic Cu(I) homeostasis. In mammals, glia, particularly astrocytes(49), have been identified as a major source of brain Cu homeostasis, with the liver primarily serving this role in the periphery. The systemic contribution of glial *swip-10* to whole body Cu(I) homeostasis parallels findings of an impact on systemic proteostasis by glia(23, 24), and can be linked to our prior demonstration of systemic oxidative stress in *swip-10* mutants(15). Although we did not examine Cu(II) levels, we hypothesize that due to loss of *swip-10* mediated Cu(II) reduction, worms will display an increase in Cu(II) due to the lack of a significant change in total Cu, possibly contributing for some or the phenotypes documented in this study Recently, a Cu(II) specific probe, similar to CF4, has been reported(50) that we plan to utilize in future studies to explore this question.

Metabolic disruptions have been implicated in risk for virtually all neurodegenerative diseases (51). Neurons are particularly dependent on mitochondrial mechanisms of ATP synthesis(52) which sustains ion gradients, synaptic release of neurotransmitters and buffering Ca^2+^ that otherwise can activate apoptotic cascades(53), increasing attention to mitochondrial modulation for therapeutic potential(6). Our hypothesis predicts that *swip-10* mutants should display deficits in Cu(I) dependent energy production, changes that would be predicted to contribute to neuronal dysfunction and possibly neurodegeneration. In this regard, we demonstrated basal OCR deficits in *swip-10* mutants using two different methods, both yielding equivalent results with respect to basal OCR. The fact that these deficits were identified in whole worms and can be rescued by glial specific expression of WT *swip-10* confirms our conclusion drawn from CF4 studies that glial *swip-10* plays a broader role than the signaling and health of glial-ensheathed neurons. Our Seahorse respirometry studies also demonstrate that *swip-10* mutants display a WT maximal OCR as well as normal non-mitochondrial OCR, typically considered to arise from oxygenase-type enzymatic reactions. Rather, *swip-10* animals possess a significantly reduced basal OCR, with pharmacological studies indicating that the deficit arises at or before Complex IV. Notably, Complex IV or cytochrome c oxidase is a multi-protein complex that relies on Cu(I) to provide for electron transfer to oxygen, creating the electromotive forced needed by Complex V for ATP synthesis. These findings are consistent with the diminished steady-state ATP levels observed in *swip-10* mutants.

It is well established that disrupted mitochondrial ATP production can lead to elevated ROS and increased cellular oxidative stress(54). Consistent with these findings, our previous work displayed a whole animal increases expression in *swip-10* animals of the ROS sensitive reporter *gst-4*:GFP(15). Since many SODs, the major intracellular class of ROS handling proteins, are Cu(I) dependent, we pursued a more direct measure of ROS in *swip-10* mutants, taking two separate approaches. First, we stained WT and *swip-10* mutants with the fluorescent ROS reporter, DCFDA observing a significant increase in whole body ROS *in vivo*. As with mitochondrial function, we achieved a normalization of whole-body ROS levels via glial expression of WT *swip-10*. Secondly, we detected diminished levels of whole body GSH that translates into a significant reduction in Redox Potential, indicative of a reduced capacity of cells, systemically, to eliminate ROS generated during normal metabolic reactions. Interestingly, we detected elevated expression of *sod-2*, a Cu(I) dependent, mitochondrial SOD (**Fig. 4**), though we observed no changes in *sod-1*, which encodes a Cu(I) independent SOD (data not shown), supporting mitochondrial deficits as a primary driver of ROS in *swip-10* mutants.

The nematode model is highly amenable to pharmacological rescue experiments that can complement and extend genetic approaches. We capitalized on this capacity to determine whether a Cu(I) deficiency is necessary and/or sufficient for the generation of one or more of the *swip-10* phenotypes studied here. The demonstration that WT worms, grown on plates that have been supplemented with the Cu(I) chelator BCS, phenocopy the OCR, ROS, gene expression, and neurodegeneration phenotypes, of *swip-10* mutants, supports a conclusion of Cu(I) necessity. With the rescue of these *swip-10* phenotypes by culturing *swip-10* mutants on plates dosed with either CuCl_2_ or ES, agents known to augment Cu(I) levels in cultured cells and animals(55), we also demonstrate Cu(I) sufficiency. ES is a particularly interesting reagent as it is a chemical Cu(I) chaperone that has been found to be efficacious for treating the Cu(I) associated deficiencies observed in a Menke’s disease mouse model as well as with human carriers of disease driving *ATP7A* mutations(56). Our studies warrant consideration of elesclomol-like agents that can boost intracellular Cu(I) levels for the treatment of neurodegenerative conditions, particularly where comorbidities, genetic markers associated with diminished *Mblac1* expression and/or the metabolic sequelae of MBLAC1 reduction can be used as biomarkers to identify those most likely to benefit.

We have shown that selective glial expression of *swip-10* can rescue both Swip, DA neuron hyperexcitability, and Glu signaling-dependent neurodegeneration. Previously, we presented evidence that the degeneration of DA neurons observed in *swip-10* mutants may be a property of glial ensheathment as we found another ensheathed neuron (OLL), known to be glutamatergic, that displays degeneration in the *swip-10* background, whereas an unsheathed neuron (BAG) shows no changes(15). Interestingly, we also found that BCS treatment induces degeneration of OLL neurons whereas BAG neurons are insensitive **(Supp. Fig. 2D)**. The basis for the Cu(I) dependent viability of glial ensheathed neurons is currently unknown. Mammalian astrocytes are known to export Cu that is acquired by neurons for metabolic use(57), and though we cannot assess such a transfer as of yet, it seems reasonable to consider this a strong possibility, particularly given the systemic phenotypes induced by head localized glial cells which may also derive from a transfer of Cu(I). Such a transfer may involve vesicle stores that fuse to release free Cu(I) or Cu(I) binding proteins analogous to mammalian ceruloplasmin(58, 59) which supports export of inter-organ transfer of Cu(I) released by the liver. Interestingly, the glia surrounding degenerating neurons in *swip-10* mutants appear morphologically normal(15), suggesting that these glia make less use of the relevant Cu(I) dependent processes whose diminished function impact the viability of ensheathed neurons. Finally, Cu(I) itself need not be transferred to the neuron and be the factor whose loss produces degeneration. We have shown that the degeneration of DA neurons in *swip-10* mutants involves excess Glu signaling by Ca^2+^ permeant Glu receptors(15). Cu(I) has been found to inhibit NMDA-type Glu receptors(59) and, if deprived of normal levels of Cu(I), DA neuron Glu receptors may be overactivated by normal levels of Glu input. It is also possible that a glial impact of low Cu(I) levels, possibly arising from altered OCR and/or ROS production, is a reduction of Glu transporter expression or function needed to limit DA neuron excitation, where chronic overstimulation of Ca^2+^-dependent Glu receptors leads to neurodegeneration. Certainly, both diminished Glu clearance and reduced export of Cu(I) may occur in parallel, triggering both local and systemic changes.

In our prior studies, we demonstrated that MBLAC1 is a high-affinity and potentially exclusive target for the neuroprotective beta lactam antibiotic ceftriaxone (CEF)(60). Conspicuously, a key function of CEF is to upregulate Glu transporters. Although CEF would be expected to block MBLAC1 action, and lead to neurodegeneration rather than neuroprotection, the limited brain penetration of systemically administered CEF may lead to induction of neuroprotective programs, a feature of which is upregulation of astrocytic Glu transporters(61). Indeed, the neuroprotective actions of CEF arise through NRF2(62-64), the mammalian ortholog of *C. elegans* skn-1, which we show to be elevated in *swip-10* mutants in a Cu(I) dependent manner(**Fig. 4C**).

Given evidence of disrupted, whole worm mitochondrial function in *swip-10* mutants, we sought to profile systemic metabolome changes in these animals to determine consequent molecular alterations that might further inform how glial-dependent Cu(I) homeostasis impacts local and systemic physiology. These experiments revealed that loss of *swip-10* results in changes in pathways arising from or compensating for losses of mitochondrial metabolism. The most significant change observed in these studies was in C-21 steroid biosynthesis (**Fig 7B).** Importantly, mitochondria play a crucial role in the metabolism of steroids, many of which are exported for use in downstream biochemical reactions critical for organismal metabolism(39, 65). Moreover, the biosynthesis of steroids in mitochondria impact mitochondria directly via the production of biochemical intermediates such as NADH(66). Thus, determining whether altered steroid hormone levels are a cause or a consequence of the metabolic insults seen in *swip-10* mutants warrants further investigation.

Compared to the restricted, glial expression of *swip-10*, mammalian *MBLAC1* is expressed widely, including in the periphery, suggesting systemic consequences of the gene’s altered expression or function. In this regard, we are reminded that the risk for AD attributed to low expression of *MBLAC1* arises in AD subjects bearing CVD co-morbidity. Similar to the brain, the heart is an energetically demanding tissue, requiring a constant supply of ATP for lifelong function. MBLAC1 is expressed in other peripheral tissues as well, including liver, the main organ supporting systemic Cu(I) homeostasis, and thus it is reasonable to expect changes in liver function as a consequence of limited MBLAC1 activity. Our prior work characterizing serum metabolome changes in *Mblac1*^-/-^ mice(26) revealed multiple altered pathways suggestive of altered liver function, including a reduction in bile acid production(67). NADH and FADH_2_ are the oxidized electron carriers supporting the mitochondrial ETC at complex I and II, respectively. The ratios of these two critical energy pathway cofactors can be used to assess the metabolic state of mitochondria(68). Recently, using a metabolic imaging approach to measure the ratios of NADH to FADH_2_ in liver sections from *Mblac1*^-/-^ mice, we found a robust decrease in this measure, consistent with a conserved mitochondrial mechanism and supporting a conserved mitochondrial impact of *swip-10* and *MBLAC1* deficiency. While work on human disorders of *Mblac1* is in its infancy, the replicated findings of AD-CDV with reduced *Mblac1* expression is compelling. In this regard, reduced cortical *MBLAC1* gene expression has been associated with AD-CDV. We therefore asked whether *swip-10* mutants might effectively model the association of reduced cortical *MBLAC1* expression with increased risk for AD-CDV. The GMC101 strain expresses human Aβ_1-42_ which forms readily detectible plaques when stained with the fluorescent Congo Red derivative, X-34(43). Here, we demonstrate that a large increase in Aβ plaque accumulation arises in worms produced to express Aβ_1-42_ on a *swip-10* mutant background. These exciting findings support use of the *swip-10* strain and *Mblac1* deficient mice to model risk and resiliency mechanisms for AD as well as PD, given that DA neurons are among the classes of neurons in the worm sensitive to *swip-10* deficiency.

## Materials and Methods

All animals were grown on what OP50 bacteria and kept under standard housekeeping protocol as previously described(69). All other methods used for these studies are reported in *SI Methods*.

## Author Contributions

PR designed and performed research, analyzed data, and wrote manuscript. VK performed research and analyzed data. ZG performed research. CLG performed research, and analyzed data. AR performed research. CDM contributed new reagents. ATP contributed new reagents. XT performed research. LC analyzed data. CJC contributed new reagents. GWM designed research, provided resources, and analyzed data. AK provided resources and analyzed data. JB performed research. RDB designed research, analyzed findings, and wrote manuscript. All authors were provided with an opportunity to review and suggest edits for the manuscript.

## Competing Interest Statement

The authors have no competing interests to disclose.

## Classification

Biological Sciences, Neuroscience

## Acknowledgments

RB gratefully acknowledges financial support from Steven and Deborah Schmidt, Grant 22A01 from the Florida Department of Health, and a pilot award from the FAU Mangurian Center for Brain Health. GM and VK were supported by NIH R01 ES 023839. CJC acknowledges funding support from the National Institutes of Health. 79465 to CJC. CJC is a CIFAR fellow. We also acknowledge the generous training, advice and/or input on the manuscript by former Blakely lab members Osama Refai, Chelsea Gibson, Maureen Hahn, and Felix Mayer.

---

**Supplemental Figure 1.**
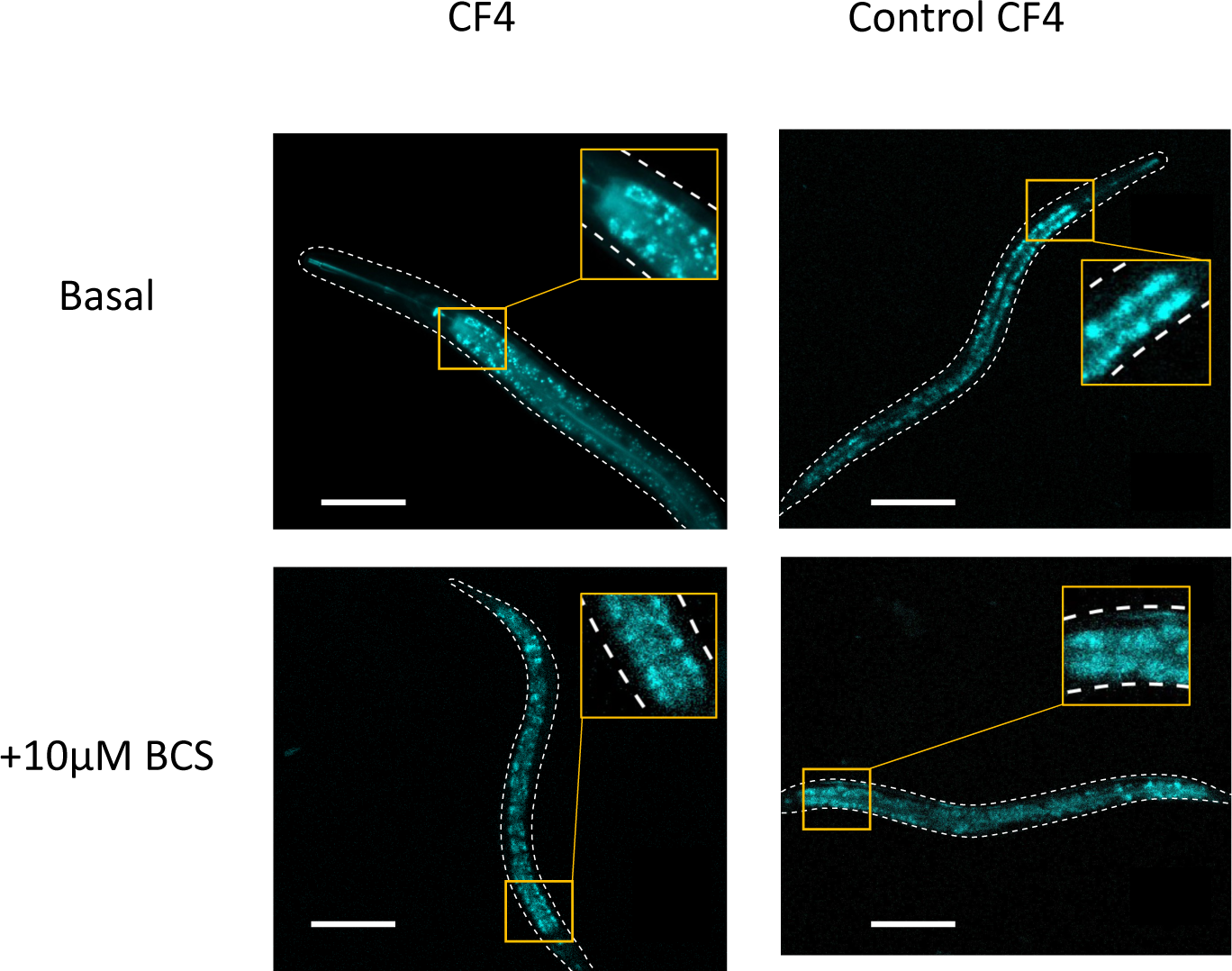
Cu(I) specificity of CF4 demonstrated by Cu(I) chelation. All images were acquired using a Nikon Confocal microscope with a 20x objective. All images were of WT (N2) animals at the L4 stage. Compared to CF4, Control CF4 has a largely reduced number and intensity of puncta. Cu(I) chelation by treating worms with 10μM BCS induced a decrease of number and intensity in puncta, similar to basal Control CF4 levels. Animals imaged with Control CF4 after being treated with BCS did not appear to have any difference in puncta count or intensity compared to basal. Scale bars equal to 50μm.

**Supplemental Figure 2.**
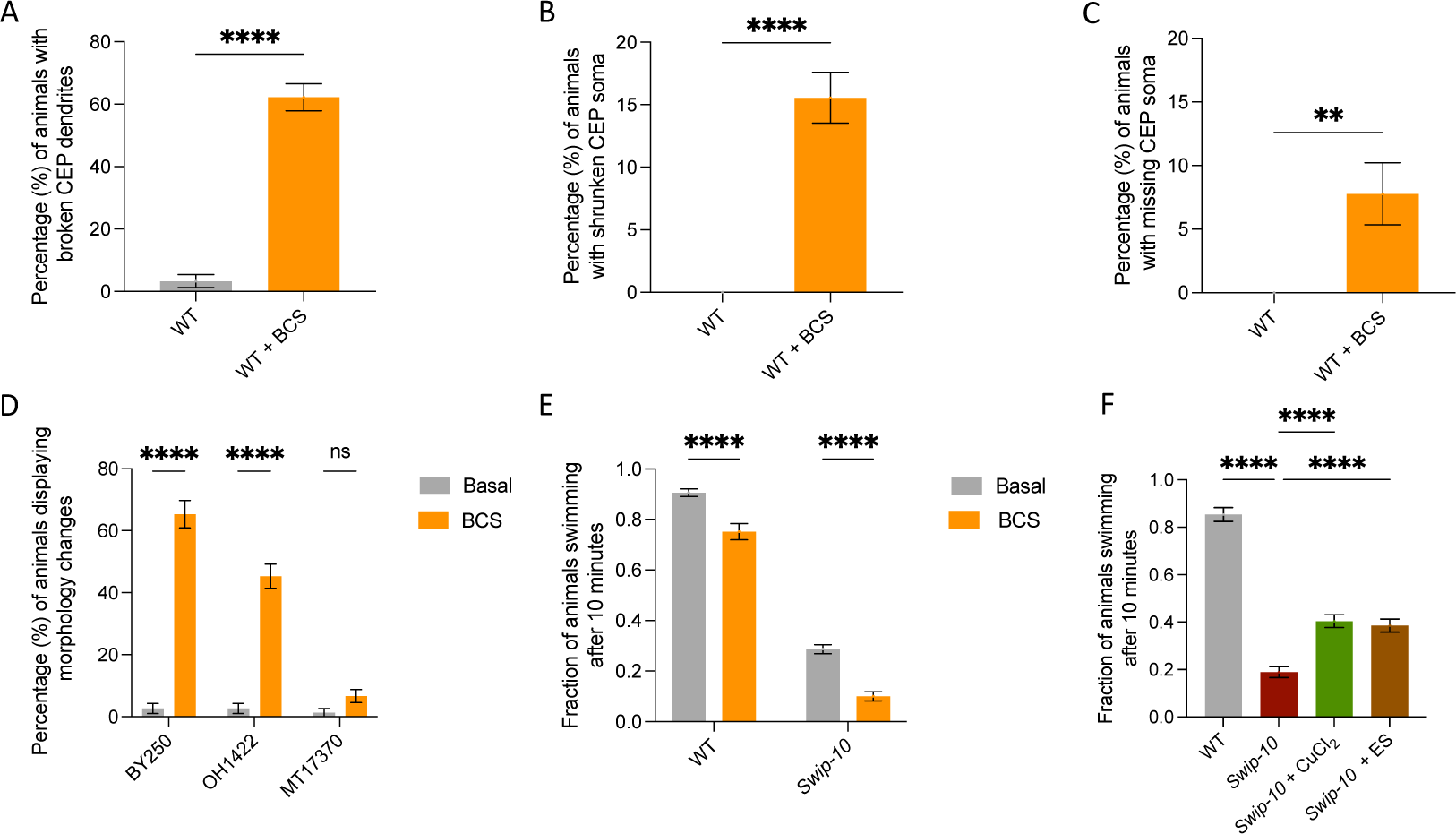
Effect of Cu^1+^ on DA neuron function and viability. A-C) DA neuron degeneration in WT [BY250(p*dat-1*::GFP)] animals treated with BCS. Statistical analyses performed using Welch’s t-test. \*\**p≤0.01, ***p*≤0.001, \*\*\*\**p*≤0.0001. 15 animals were measured in each independent replicate. N=5 for all conditions measured. All measurements were performed blinded by manually visualizing DA neurons in in-tact living animals on epifluorescence microscope, paralyzed using levamisole solution. A) Average of population percentage of animals showing a break in the continuous GFP visualization in dendrite track. B) Average of population percentage of animals displaying reduced soma size. C) Average of population percentage of animals completely missing one or more DA neuron. D) Population percentage of animals displaying one or more morphological alteration in neuronal GFP. OH1422 is a strain harboring a GFP marker in OLL neurons (ser-2(prom3)::GFP + rol-6(su1006)). MT17370 is a strain harboring a GFP marker in BAG neurons (nIs242 [Pgcy-33::gfp] III; lin-15AB(n765) X)). Statistical analyses performed using Ordinary Two-way ANOVA. \*\*\*\**p*≤0.0001. E) Testing the effect on swip of animals grown on plates containing BCS. F) Testing the effect on Swip of animals grown on plates containing either CuCl_2_ or elesclomol. E,F) Manual Swip measurements of animals placed in water. Bars displayed as fraction of animals swimming after 10 minutes after being placed in water. Ordinary One-way ANOVA.

**Supplemental Figure 3.**
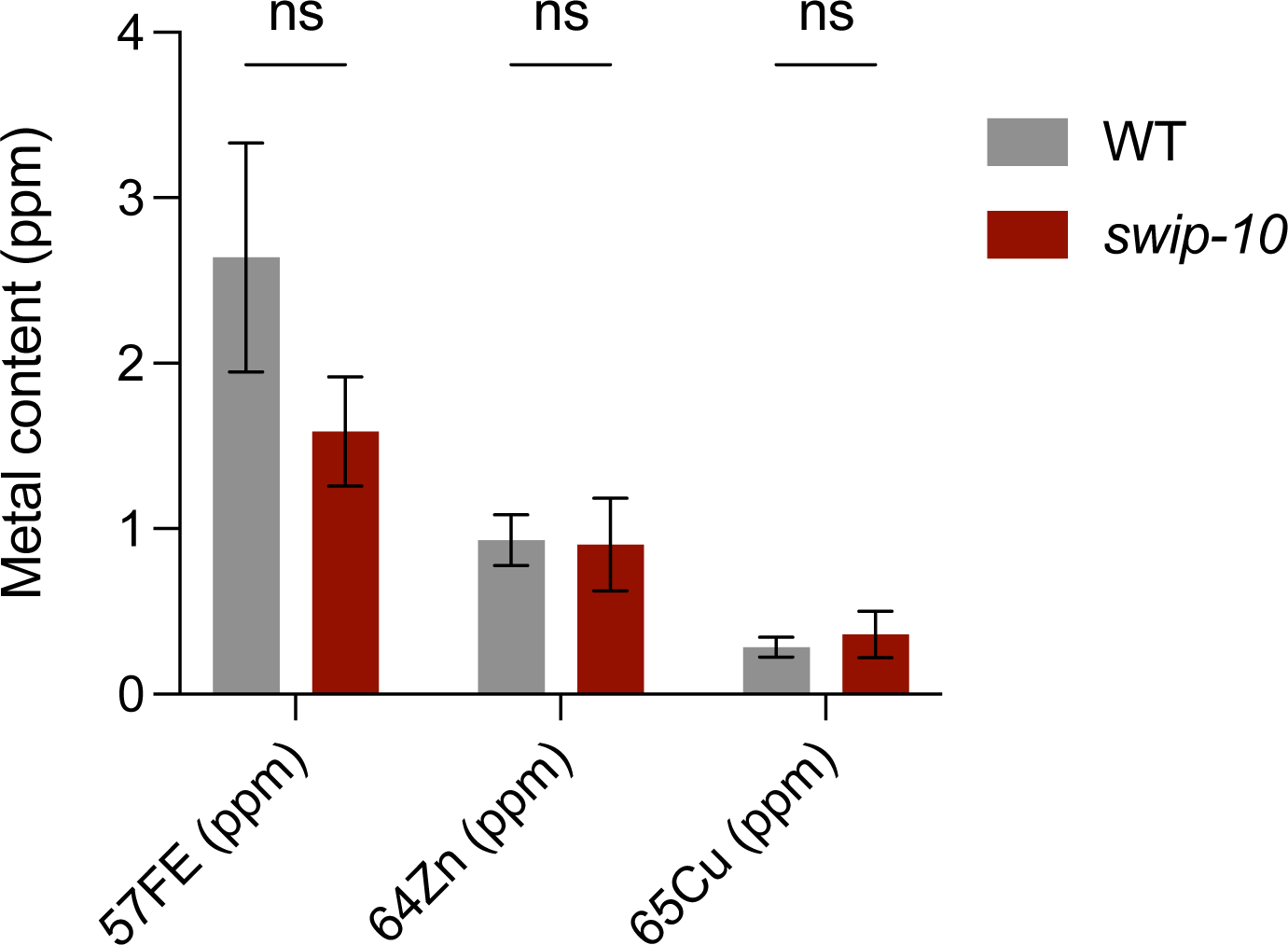
No difference in stable metal isotope concentrations between WT and *swip-10* animals. Mass spectrometry-based measurements of the most stable isotopes of iron (57FE), zinc (64Zn) and copper (65Cu). Measurements were from whole worm content and reveal no significant differences between genotypes. Statistical analysis performed using two-way ANOVA, with a Sidak’s multiple comparisons test. ns, non-significant.

**Table 1.**
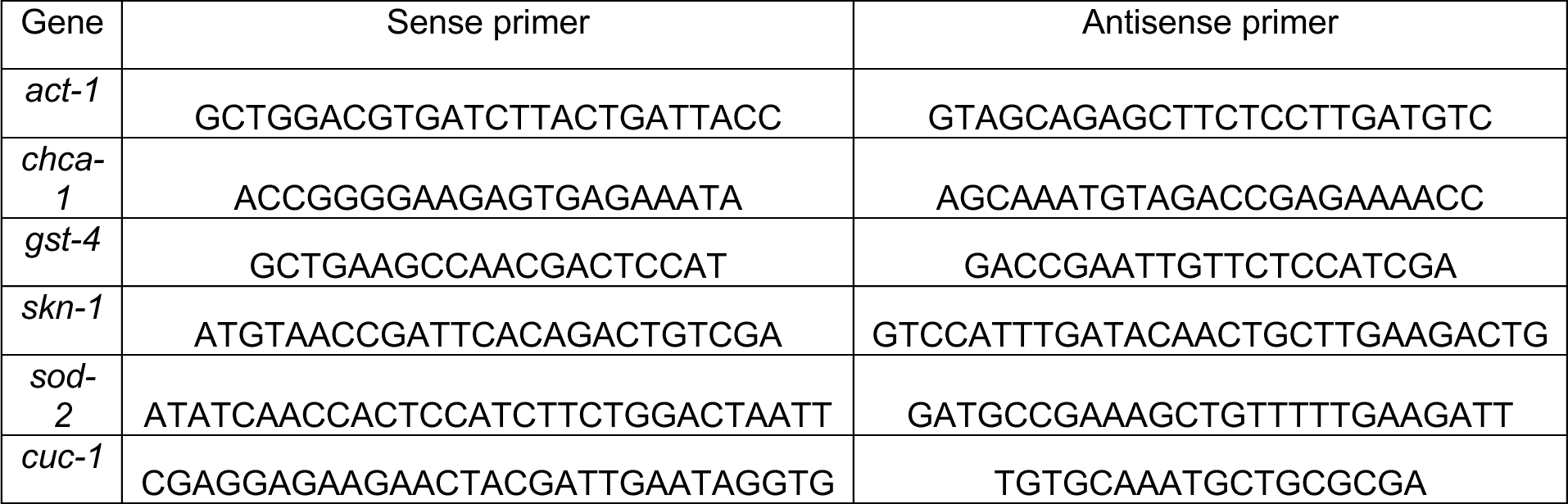
Oligonucleotide primer sequences used for quantitative PCR analysis of mRNA expression.

## Notes

### Competing Interest Statement

The authors have declared no competing interest.

