## Supplementary material for "*Glial swip-10* expression controls systemic mitochondrial function, oxidative stress, and neuronal viability via copper ion homeostasis": SI Methods

### Materials and Methods

*C.elegans* strains and husbandry

All animals were grown on what OP50 bacteria and kept under standard housekeeping protocol as previously described(66). N2(Bristol), our WT strain, and the animals harboring a tm5915 deletion allele of the *swip-10* gene, were used for all experiments. The BY1199 strain [vtIs7;*swip-10*(tm5915); Ex269 (P*ptr-10*::*swip-10*cDNA::GFP, P*unc-122*::RFP)] was used for driving glial specific expression of *swip-10*. The GMC101 strain(41) was used to investigate plaque formation by Ab$_{1-42}$ in the context of a *swip-10* mutation to model the risk of reduced MBLAC1 expression associated with AD-CDV. To visualize CEP DA neurons, the BY250 [P*dat-1*::GFP] strain was used. To visualize OLL neurons the OH1422 [otIs138 (ser2prom3::GFP + rol-6) X] strain was used. To visualize BAG neurons the MT17370 [lin-15AB(n765);nIs242[Pgcy-33::GFP]] strain was used. 2′,7′-Dichlorofluorescin diacetate (DCFDA) was obtained from Millipore Sigma. Bathocuproinedisulfonic acid, disodium salt hydrate (BCS), 97% was obtained from Fischer Scientific. All BCS stocks were solubilized in MilliQ water. Elesclomol (ES) was obtained from Selleckchem, solubilized in DMSO to make stocks of 10mM. $CuCl_2$ was obtained from Sigma-Aldrich, dissolved in MilliQ water to obtain stocks of 10mM. Lipophilic Congo red derivative, X-34 was obtained from Millipore Sigma. X-34 was initially solubilized in DMSO, further diluted for worm staining using a 10mM solution of Tris. CF4 and CF4 control Cu(I) probes were synthesized by the C. J. Chang laboratory at UC Berkeley.

Cu(I) measurements

As described previously(29), CF4 can be used to stain specifically for the presence of Cu(I) in *C. elegans in vivo*. In our studies, L4 stage animals were treated for 12-16 hours with a 25μM CF4 solution prior to confocal imaging. 25μM working solution was prepared from a 5mM stock (DMSO as solvent) by dilution in M9 buffer. Z-stack images were obtained on a Nikon A1R confocal microscope for each animal to capture all focal planes showing intestinal puncta. Z-stack images were then collapsed into maximum intensity projections. For quantitation, the maximum intensity images were used to manually count CF4-labeled puncta. All experiments involving imaging, quantitation and analysis were performed blinded to genotype. A total of five animals were quantified over five separate days for each condition. Statistical analyses performed using Student's T-test. ****p* ≤0.001.

Oxygen consumption rate (OCR) assays

High-resolution respirometry experiments were performed on whole animals using the Oroboros Oxygraph 2k respirometer. Oxygen flux was recorded in real-time at 20°C under constant stirring at 300 rpm in M9 media(67). Synchronized worms were collected at the L4 stage for all experiments noted. Animals were washed off agar plates using M9 buffer to a total volume in a conical tube of 10ml. These tubes containing worms and M9 were centrifuged at 1200xrpm for 1 min to collect a pellet of worms from which the supernatant was aspirated. Washing via centrifugation was performed 3X before all experiments. After washing animals, the amount of liquid in each tube was reduced to approximately 4mL. 20μL of the worm suspension was placed on a slide to visually count number of worms per microliter. To obtain approximately two animals per microliter, dilutions were performed as necessary. Once the solution was at a volume where it contained approximately one worm per microliter, these animals were transferred into the Oroboros Oxygraph 2k respirometer. Using the solution containing one worm per microliter, 400μL was added to the respirometer chamber for a total of 400 animals per chamber. Oxygen flux during a stable 10 minute window was analyzed offline by Datlab software (Oroboros

**Commented [RB1]:** My understanding is that we were going to give a small amount of info in Methods and then refer the reader to the Supplement for additional description. This would then be considered Supp 1. Here the Methods are not in the Supp.

**Commented [AK2]:** In Figures 2, 4 and 5, the O2k measurements have pmol/s for the units. I'm wondering what the best approach would be, since we encountered this expressed in the literature in different ways.
I'm assuming this is representing pmol/s in 400 worms? If so, we may need to consider mentioning this directly, probably near the end of this O2k methods section.

**Commented [AK3]:** Reference below if needed. This was one of the papers we used early on when we were planning and tinkering. I can pull others as well, please let me know, Constitutive MAP-kinase activation suppresses germline apoptosis in NTH-1 DNA glycosylase deficient C. elegans - PubMed (nih.gov)

Instruments, Innsbruck, AT). To determine the basal OCR, measurements from each biological condition were averaged from two technical replicates in different chambers of the same machine.

Alternatively, whole-worm respiration was measured as a function of time using a Seahorse XF24 analyzer (Agilent) and with methods previously described(30). Briefly, worms were plated in M9 buffer. M9 buffer without any worms was used as the blank. Baseline respiration was measured, followed by injection of FCCP (10 µM, final concentration) to elicit maximal respiration, followed by sodium azide (40 mM, final concentration) to account for non-mitochondrial respiration. A total of 20 worms per well were used for recordings. Measurements taken every 2 min were normalized to number of worms per well.

<u>ATP assays</u>

All assessments of ATP levels were obtained with L4 animals. Animals were washed off plates using M9 buffer and collected into 15mL polystyrene conical tubes, then washed 3X via centrifugation, and then transferred to 1.5mL microcentrifuge tubes. Final volume in microcentrifuge tubes was 1.0mL. Tubes containing worms and M9 were subjected to 5 cycles of freezing in liquid nitrogen and subsequent rapid thawing in boiling water for tissue dissociation. Protein content of samples was determined via a BCA protein assay kit (ThermoFisher).Each sample was subsequently resuspended to 2µg/µL by dilution with M9. Subsequently, 90µL of luciferase reagent (ATP determination kit, ThermoFisher) was added to the wells of an opaque 96-well plate at room temperature followed by addition of 10µL of either protein standard or sample. In a light shielded environment, sample plates were incubated at $37^{o}$C for 10 min. ATP-dependent luciferase activity was measured using a Microbeta 3 plate reader (Perkin Elmer). Absolute ATP concentrations of samples were determined using an ATP standard curve.

<u>ROS quantification using DCFDA</u>

ROS measurements were performed using 2',7'-Dichlorofluorescin diacetate (DCFDA) as described previously(68). Briefly, animals were collected in M9 buffer, washed 3X before treatment for 1hr at room temp in a 50µM DCFDA solution. DCFDA solution was prepared in DMSO. Solutions and treated worms were shielded from light before microscopy. After 1hr, animals were washed in M9, and transferred to agar plates. Confocal microscopy was performed to acquire images of DCFDA treated animals. A 488nm wavelength laser was used for excitation. Analyses were performed by acquiring images at maximum pixel intensity, divided by the area of the animal ($mm^2$). Ten animals (per condition) were imaged to average values. Data were collected from five subsequent experiments. Data are represented as fold change relative to N2 animals.

<u>Reduced and oxidized glutathione assays</u>

Measurement of reduced and oxidized glutathione was pursued using a protocol described by Jones et al (1998), adapted for *C. elegans*. Here, 500 worms per strain were collected using a Union Biometrica COPAS Biosorter (Holliston, MA) into 500µL of preservation buffer (5% (w/v) perchloric acid containing 0.2 M boric acid, and 10mM γ-glutamylglutamate (γ-Glu-Glu)). Samples were snap frozen in liquid nitrogen and stored at -80°C. On the day of analysis, samples were allowed to thaw, and sonicated on ice using 12 pulses at 50% amplitude for 5 secs with a 2 sec break. Samples were centrifuged at 13,000 rpm for 2 min, with 300µL of the supernatant then transferred to a 1.5mL microfuge tube and processed as described by Jones et al. Briefly, 60µL of iodoacetic acid solution (40mM) was added to each tube. A potassium hydroxide (KOH)/tetrahydroborate solution was used for adjustment of pH to 9.0 ± 0.2. After 20 min of incubation at room temp, 300µL of 20 mg/ml dansyl chloride solution in acetone was added.

**Commented [KV4]:** The worms were sorted into this buffer already present in the collection tube. This made sure they were preserved as soon as they were sorted.

Samples were vortexed and placed in the dark for 24 hrs. Subsequently, 500μL of chloroform was added to each sample and vortexed, followed by centrifugation at 10,000 rpm for 2 min to separate organic and aqueous phases. The upper aqueous layer was collected into HPLC vials with 20μL of each sample injected onto a 3-aminopropyl column. All analysis parameters were as previously described(69). Quantitation of reduced and oxidized glutathione were obtained by peak integration relative to the internal standard.

RT-qPCR based gene expression analyses

*Swip-10* mutants and WT worms were grown on NGM plates until the L4 stage when they were collected for whole animal RNA isolation using a rapid, proteinase K-based protocol outlined as previously described(70). cDNA libraries were prepared with Maxima H Minus First Strand cDNA synthesis Kit (catalog no. K1652, ThermoFisher) using the whole animal RNA isolate as template material. Before cDNA library preparation, all RNA samples were treated with Turbo DNA-free kit (catalog no. AM1907, ThermoFisher) to eliminate any potential genomic DNA. cDNA libraries generated were then used for targeted quantification of gene expression using the PowerUp SYBR Green Master Mix (Catalog no. A25780, ThermoFisher). Oligonucleotide primers used to amplify cDNA in qPCR reactions are listed in Table 1. All data derive from at least 6 biological replicates of prepared RNA templates. Two technical replicates were used for each sample. For each gene quantified, the fold changes in expression levels of mutants were normalized to WT levels using Actin (*act-1*).

**Commented [RB5]:** How many technical replicates

ICP-MS assays for levels of Cu and other divalent metals

To measure total concentrations of various metals, liquid ICP-MC analyses using L4 stage worms were carried out as described previously(28). Briefly, worms were washed 3X with M9 and then dropped using a glass pipette into liquid nitrogen to make small frozen pellets. Approximately 100mg of these pellets were collected and frozen for subsequent analysis. Samples were digested in 1 mL trace-metals grade concentrated nitric acid (BDH Aristar Ultra, 87003-228) per 100 mg of tissue at room temp overnight, then diluted in 2% nitric acid with Gallium as internal standard. ICP-MS analyses were performed with an iCAP-Qc ICP-MS (ThermoFischer).

Imaging and quantification of neurodegeneration

Quantification of the degeneration of *C. elegans* cephalic (CEP) DA neurons, OLL neurons and BAG neurons was performed as outlined previously(15). Briefly, all experiments were performed on young adult animals, one day after the L4 stage. Strain BY250 [P*dat-1*::GFP (vtIs7)] was used for all experiments. Neurodegeneration was scored using a Zeiss upright compound epifluorescence microscope (20x objective). Truncated dendritic processes, shrunken soma and missing neurons were measured as outlined previously(15). 15 animals were scored each day for at least 5-7 days for each condition. All experiments and analyses were performed blinded to genotype.

Swip assays

Swimming Induced Paralysis (Swip) assays performed here have been described previously(12). All experiments noted in this study were performed using L4 stage animals with manual counting of number of animals paralyzed 10 minutes after submerging in distilled $H_2O$. 10 animals were placed into wells containing 100μL of fresh milliQ water, each well consists of one biological sample. A minimum of 32 wells were assessed for each condition. Analysis was carried out over at least 3 subsequent days to account for variability in culturing of worms.

Metabolomics analyses by liquid chromatography coupled to high-resolution mass spectrometry (LC-HRMS)

**Commented [GM6]:** The language had evolved- high-resolution mass spectrometry (LC-HRMS) is best

*Swip-10* mutants and WT worms were grown on NGM plates until the L4 stage. Worms were then washed off plates using M9 and sorted into five replicates with 500 worms each, for each strain, using a Union Biometrica COPAS Biosorter (Holliston, MA), snap frozen in liquid nitrogen, and stored at -80°C until needed for processing. Metabolites were extracted using acetonitrile (in a 2:1 ratio). The extraction solvent also contained internal standards. Bead-beating was used to disrupt the worm cuticle and improve extraction. A spatula-full (~50 beads) of acid-washed beads (300 microns) were added to each tube. Each sample was placed in the bead beater at speed 6.5 m/s for 30 seconds, allowed to equilibrate on ice for one minute, and placed in the beater for another 30 seconds, at the same speed. All processing was performed on ice or in a cold room when necessary. Untargeted high-resolution mass spectrometry was performed at the Clinical Biomarkers lab at Emory University using a HILIC column (positive ionization) and C18 column (negative ionization) using chromatographic methods previously described (Liu et al. 2016). Mass spectral data was generated on a Thermo Orbitrap Velos in full scan mode, scanning for mass range 85 to 1250 Da. Data were extracted using the R packages apLCMS (Yu et al. 2009) and xMSanalyzer (Uppal et al. 2013). The HILIC column detected 20,672 features which reduced to 5,927 when filtered using features detected from gut clearance. The C18 column measured 12,039 features which reduced to 2,960 when filtered. A feature was retained if its intensity was more than twice that of gut clearance in the *swip-10* mutant samples. Features were imputed if missing with half the value of the minimum abundance, log10 transformed, and auto scaled. All data processing, analysis and visualization was done in R (version 3.6.0) and Metaboanalyst(35). Pathway analysis was done using the mummichog algorithm version 2 (Li et al. 2013) using the human metabolic reference map.  A p-value cut-off of 0.01 was used to identify pathways of interest altered in the *swip-10* mutant using features measured on the HILIC positive column. A p-value cut-off of 0.05 was used for pathway analysis using features measured on the C18 negative column.

Quantification of $A\beta_{1-42}$ plaques

L4 stage animals were utilized for all experiments. Animals were stained with X-34, as described previously(41). After staining for 2hrs, animals were washed in M9 buffer 3X, then transferred to plates for 6 hours, shielded from light. The animals were then imaged using a Keyence BZ-X series microscope using a 20x objective and a DAPI fluorescent filter. Plaques that appeared in the head region, anterior of the pharyngeal bulb were manually counted.

Statistics and graphics generation

All analyses and figure curation were carried out in Graphpad Prism version 10. All experiments and analyses were performed blinded to genotype. Fluorescence intensity measurements were performed using FUJI (ImageJ). For all analyses, a *P*<.05 was considered statistically significant. Tests of statistical significance are provided in figure legends.

**Commented [RB7]:** Is your version Version 10? Mine is version 8. Perhaps I just didn’t keep my own Prism as updated as the lab’s or yours?
